# Leveraging a Billion-Edge Knowledge Graph for Drug Re-purposing and Target Prioritization using Genomically-Informed Subgraphs

**DOI:** 10.1101/2022.12.20.521235

**Authors:** Brian Martin, Howard J. Jacob, Philip Hajduk, Elaine Wolfe, Loren Chen, Henry Crosby, Matthew Lefever, Richard Wendell

## Abstract

Drug development is a resource and time-intensive process resulting in attrition rates of up to 90%. As a result, repurposing existing drugs with established safety and pharmacokinetic profiles is gaining traction as a way of accelerating therapeutics development. Here we have developed unique machine learning-driven Natural Language Processing and biomedical semantic technologies that mine over 53 million biomedical documents to automate the generation of a 911M edge knowledge graph. We then applied subgraph queries that relate drugs to diseases using genetic evidence to identify potential drug repurposing candidates for a broad range of diseases. We use Carney Complex, a disease with no known treatment, to illustrate our approach. This analysis revealed Ruxolitinib (Incyte, trade name Jakafi), a JAK1/2 inhibitor with an established safety and efficacy profile approved to treat myelofibrosis, as a potential candidate for the treatment of Carney Complex through off-target drug activity.

## Introduction

The research and development cycle of a novel drug is extensive, requiring an estimated 1-2 billion dollars of investment on average and 10-15 years to reach the market from conception through FDA approval.^1,2^ Ultimately, only an estimated 10-15% of drugs from phase I of clinical trials are approved for market use.^3^ As such, drug repurposing methods to find new applications for currently approved or drugs that failed their initial intended purpose is gaining increasing interest as a method for accelerating therapeutics development.^4^ As existing on-market drugs have established safety and pharmacokinetic profiles, drug repurposing can shorten the development cycle and mitigate risks early in the process.^5^ An extreme example of successfully repurposed drugs is thalidomide. Originally marketed as a morning sickness drug that was pulled from the European market for severe birth defects in the 1960s, it was ultimately brought back as a treatment for a range of diseases, including many forms of cancer and immune diseases.^6^

Leveraging large, complex drug-disease networks has emerged as a powerful approach for indication expansion and drug repurposing, particularly for identifying existing therapeutics that can treat rare, intractable, or emerging (i.e. viral) diseases.^7–10^ In network-based approaches, relationships can be inferred between entities such as drug-gene or drug-disease as entities within the same biological network.^11^ While graph and network approaches are not novel, new strategies have been developed in recent years to execute these queries in a systematic and rational way. Natural Language Processing (NLP) of biomedical literature is a rapidly maturing approach to identifying biological entities and extracting relations and insights from structured and unstructured text.^12^ As 80% of medical data is unstructured, using NLP to extract potential associations between chemicals/drugs, target genes and proteins, and disease states from biomedical text provides a powerful avenue for accelerating drug discovery and repurposing.^13^ In addition, as previous work has demonstrated a significant increase in odds ratio (>2X) for gaining drug approval when genetic evidence links a gene and a targeted disease,^14,15^ there is great potential to integrate NLP-derived drug-disease associations with available genetic and genomic data to focus on genetically supported drug repurposing.

Here, we describe the results of a subgraph pattern matching methodology, along with potential applications leveraging a multimodal knowledge graph built from various public data sources (both structured and unstructured) that have been harmonized to common ontologies. We demonstrate that graph-based inferences of drug-disease associations are significantly enriched by requiring direct or indirect (i.e., pathway-based) genomic evidence within the subgraph. We also document that hundreds of thousands of genomically-informed subgraphs can be identified that associate a known drug with a novel disease. We hypothesized that these novel drug-disease associations could be used to evaluate potential drug re-purposing opportunities, and we tested this hypothesis on Carney Complex - a rare disease with no available treatments.

## Methods

### Construction of the Knowledge Graph

Our knowledge graph network consists of 911 million edges and enables the development of graph queries that identify specific subgraph patterns between start and end nodes (i.e., a drug and a disease) and can be calculated at scale. Generating the knowledge graph involved automating large-scale curation of numerous unstructured and structured data sources, integrating the resulting metadata into a hybrid index of databases spanning graph, text and structured databases, and deploying graph analytic models that reduce knowledge graph data into subgraphs.

The curation of unstructured data was accomplished by deploying proprietary machine learning algorithms that perform four specific tasks: Named Entity Recognition (NER), Biomedical Ambiguity Resolution (BAR), Semantic Relationship Quantification (SRQ) and Causal Context Expression Curation (CEC). NER models include deep neural network models within spaCy frameworks that are custom trained to recognize drug discovery domain entities such as genes, drugs, disorders, phenotypes, adverse events, etc. The BAR processes employ custom machine learning algorithms that resolve synonyms, acronyms and ambiguous terms to enhanced versions of biomedical ontologies (e.g., HGNC, EFO, MESH, etc.). The base model for SRQ was a large transformer-style model pre-trained on a large multi-domain corpus of scientific text and fine-tuned on expertly annotated gold standard data. The SRQ model outputs a score representing the confidence of a meaningful semantic relationship between the two entities in a sentence. All resulting candidate relationships evaluated by the SRQ model are given a semantic relationship prediction score between 0 and 1. A score of 0 represents a prediction that entities mentioned within a single sentence merely co-occur and do not have a meaningful semantic relationship, while a score approaching 1 indicates a prediction of a high likelihood that two biomedical entities are semantically related. For example, consider the sentence in Figure 1. In this figure, three gene entities were extracted and mapped by the NER-BAR pipeline components. The pairwise combination of these extractions results in three candidate relationships to be evaluated by the SQR model. In this example, the model evaluates that this sentence does not by itself represent evidence of a semantic relationship between **TRAF6** and **PLK**, returning an SQR score of 0.32. However, it does represent evidence of a relationship between each of those and **KLF4** with the model returning a score of 0.86 between **TRAF6** and **KLF4** and 0.90 between **PLK** and **KLF4**. This differs from other approaches that may only evaluate relationships simply by syntactic proximity, which correlates fewer words between entities to stronger semantic relationships, or by co-occurrence in a sentence or document. Raw semantic scores from the SRQ model were then rescaled such that a threshold of 0.5 maximized model performance against the validation set of annotated data. Relationships with SQR scores of below 0.5 were filtered out. Those with SQR scores of 0.5 and greater were used to construct the knowledge graph, and semantic scores were rescaled with an affine transformation from 0.5-1 to 0-1, rounding to the nearest 0.2 cutoffs. The end value is then presented as the “confidence” of a single piece of evidence. The CEC, built using syntactic dependency graphs, then processes the directionality of entities in cause-and-effect relationships, along with biomedical verbs and adverbs that are foundational for scientific insight. The curation of structured data sources involved transforming source fields to align with the same data schema outputted by the unstructured processes noted above.

**Figure 1.**
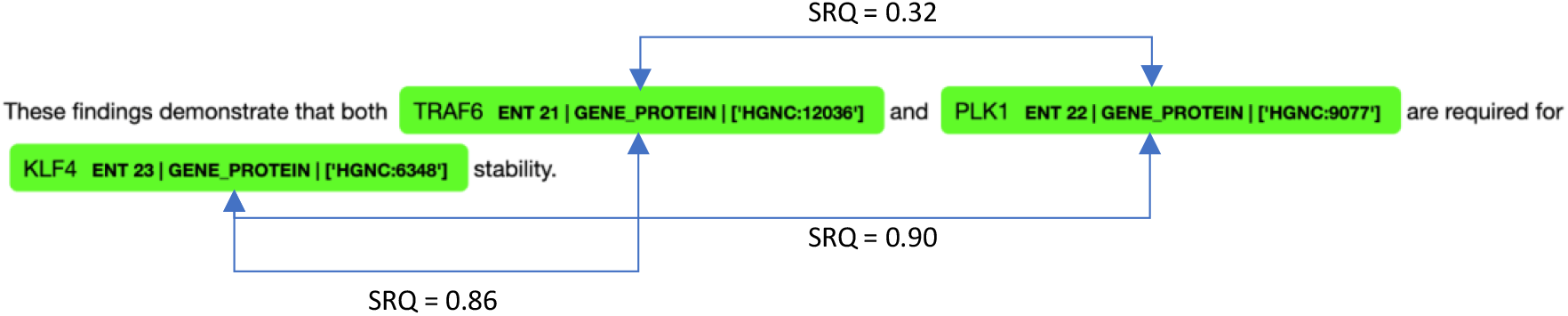
Example of Named Entity Recognition (NER) and Biomedical Ambiguity Resolution (BAR) processes applied to a sentence from the biomedical literature (tagging the terms highlighted in green), with resulting Semantic Relationship Quantification (SRQ) scores as described in the text.

The resulting knowledge graph is then stored in a database architecture that employed a graph database for graph data, a SOLR database for text format data and BigQuery for structured data. All data across the databases are interconnected with metadata links that enable the analysis to span graph format, text format and structured format data. This database architecture enables insights from subgraphs generated by the graph databases paper to index source text from source literature and patent documents as well as compare subgraph patterns to structured data from clinical trials.

While publicly available data sources were utilized in constructing the tellic knowledge graph used in this analysis, customer-specific deployments that include proprietary data have also been used to validate the conclusions. Public data sources typically fall into two categories: unstructured text from PubMed, PMC, bioRxiv, medRxiv, arXiv, Google Patents and NIH grants, and structured data with a standardized format (e.g., HGNC, GWAS, dbSNP, and CPDB).

### Subgraph Identification and Validation

To assess the inference power of these sub-graphs, a validation set of 96,707 clinically supported Drug-Disease relationships was defined, derived from a set of 6,226 drugs tested in Phase 1-4 clinical studies against 2,840 unique indications. These data are supplied as Supplementary Material.

### Drug-Disease Subgraph Models

With a graph containing over 900 million relationships, a large number of subgraphs can be created to answer a wide range of questions. For this paper, we have chosen relatively simple subgraphs that relate a Drug to a Disease through a specific gene or genes, with or without genetic evidence, as shown in Figure 1. Such subgraphs can be used to assess the potential for a given drug to have therapeutic value in the treatment of an associated Disease. The Gene-Disease relationship was defined in three different graph representations requiring either: 1) direct literature support between both a drug and a gene and that same gene and a disease, resulting in a subgraph that contains only three entities (DD3, Figure 1A), 2) direct literature support between a drug and a gene and a known variant of that gene that has an association with a disease, resulting in a subgraph of four entities (DD4, Figure 1B), or 3) direct literature support between a drug and a gene, a gene1-gene2 relationship based on known pathways, and known variant of the second gene that has an association with a disease. This results in a subgraph containing five linked entities (DD5, Figure 1C). For this analysis, pathway membership for DD5 was defined as all gene pairs with confidence scores > 0.8 in the Consensus Path Database (CPDB). All relationships extracted from unstructured data required a minimum of 5 pieces of evidence and an average semantic confidence score of >0.5 across all evidence. A list of source data for each relationship type is included in Supplemental Materials.

## Results and Discussion

The total counts of all DD3, DD4, and DD5 subgraphs identified in the tellic graph are shown in the second column in Table 1 (column titled ‘Total Subgraph Counts'). For purposes of validating the DD3, DD4 and DD5 subgraphs, a subset of known drugs with reported clinical trial activity against known diseases was constructed, such that the enrichment of clinically supported Drug-Disease associations using the subgraphs described in this work could be assessed. A set of 6,226 drugs that have achieved at least Phase 1 status were identified, which were reported in clinical trials for a set of 2,840 unique diseases (see Supplemental Material). This results in more than 17 million random pairwise connections between all drugs and diseases in this set, while only 96.7K drug-disease pairs are reported to have clinical trial activity. The validation set for the DD3, DD4, and DD5 subgraphs was then limited to these specific Drugs and Diseases, with the resulting counts shown in column 3 of Table 1 (column titled ‘Drug-Indication Subgraph Counts’). From these data, the ‘random’ chance of a drug-disease pair having clinical support is 0.55% (96.7K clinically supported out of 17 million total possible pairs). In the 37M DD3 subgraphs, there are 4.7M unique drug-indication pairs in the validation set, 73K of which are clinically supported, resulting in a validation ratio of 1.56%. While this 2.85-fold enrichment over random suggests that the availability of literature support increases the likelihood of a drug-disease association having clinical support, it is likely biased by the fact that nearly all drug-disease and gene-disease pairs being tested in clinical trials would be supported by literature evidence. Requiring at least one genetic association between the gene and disease significantly decreases the overall subgraph count but increases the validation ratio to 8.76% (DD4-1). This validation ratio increases to 19.52% when at least five variants (DD4-5) that mediate a gene-disease association have been reported, indicating that increasing levels of genetic support leads to increasing levels of clinically supported drug-disease associations. These levels of enrichment were observed even when the clinical support was required to be Phase 2 and beyond and across oncology and non-oncology indications (see Supplemental Material). Note that while all variants in DD4-2 through DD4-5 were associated with the same gene and indication, no attempt was made to cluster variants based on their chromosomal location. A similar pattern could be observed within the DD5 subgraphs (see bottom 5 rows in Table 1), which add an additional connection between genes sharing the same pathway. The overall enrichment values are lower than those of DD4 subgraphs, likely due to the subgraph pattern resulting in a larger count of subgraphs and a subsequent “dilution” of the data. However, the overall pattern of an increasing validation ratio corresponding to the level of genetic support holds, also suggestive of more novel opportunities for drug repurposing being found within this analysis pattern.

**Table 1.** Subgraph pattern count of DD3, DD4 and DD5 with clinical evidence enrichment based on semantic relationships

| Subgraph Pattern | Total Subgraph Counts | Drug-Indication Subgraph Counts | Documented Phase 1+ | Clinically Supported | Enrichment |
| --- | --- | --- | --- | --- | --- |
| All pairs | 17,681,840 | 17,681,840 | 96,707 | 0.55% | - |
| DD3 | 37,304,935 | 4,708,463 | 73,425 | 1.56% | 2.85 |
| DD4-1 ($\geq 1$ Variant) | 517,044 | 219,000 | 19,185 | 8.76% | 16.02 |
| DD4-2 ($\geq 2$ Variant) | 181,916 | 93,212 | 11,893 | 12.76% | 23.33 |
| DD4-3 ($\geq 3$ Variant) | 80,480 | 49,095 | 7,852 | 15.99% | 29.24 |
| DD4-4 ($\geq 4$ Variant) | 48,796 | 32,100 | 5,563 | 17.33% | 31.69 |
| DD4-5 ($\geq 5$ Variant) | 31,701 | 21,487 | 4,195 | 19.52% | 35.70 |
| DD5-1 ($\geq 1$ Variant) | 4,467,507 | 770,519 | 31,302 | 4.06 | 7.43 |
| DD5-2 ($\geq 2$ Variant) | 1,730,087 | 376,583 | 21,448 | 5.70 | 10.41 |
| DD5-3 ($\geq 3$ Variant) | 948,198 | 240,034 | 16,275 | 6.78 | 12.40 |
| DD5-4 ($\geq 4$ Variant) | 668,278 | 179,316 | 12,893 | 7.19 | 13.15 |
| DD5-5 ($\geq 5$ Variant) | 523,476 | 142,842 | 10,712 | 7.50 | 13.71 |

This work defines two subgraph patterns (DD4 and DD5) that algorithmically encode such genetic associations and can be routinely applied at scale across the constructed knowledge graph. The resulting inferences can now be further investigated to identify new targets with some level of drug validation or new therapeutic opportunities for existing drugs.

### Application to Rare Genetic Diseases

The DD4 and DD5 sub-graphs described in this work can be used to complement target validation activities, looking for genes or gene pathways that have both a genetic association to a disease and some evidence of clinical activity with at least early stage (Phase 1 and beyond) drug candidates. When limited to on-market drugs, these sub-graphs then represent potential drug-repurposing opportunities. This can be useful for indication expansion, where the genetic evidence supports treatment of a disease that is pathophysiologically related to an existing indication for that drug. When limited to on-market drugs, such graphs also represent an opportunity to identify potential novel treatments for rare diseases.

There is an urgent need to find on-market drugs to treat rare diseases. Unfortunately, clinical trials for rare diseases may be too slow to help patients and too costly to justify the investment by most pharmaceutical companies. Thus, repurposing on-market drugs gives the physician and patients a potential avenue for treatment. Multiple approaches to repurposing drugs for the treatment of rare diseases have led to the identification of promising candidate drugs, some of which are in the advanced stages of clinical trials or already approved.^4,5^ Despite these efforts, it is estimated that fewer than 6% of all rare diseases have an approved treatment option, highlighting the tremendous unmet medical need.^16^ As the specific gene or genes involved in the onset and progression of these diseases are known, the DD4 and DD5 as subgraphs described here can be restricted to include at least one of these genes, yielding a focused number of genetically supported drug-disease inferences that can be further evaluated.

We demonstrate the use of this knowledge graph for the identification of potential drug-repurposing candidates against Carney Complex (CNC). CNC is a rare multiple endocrine neoplasia (MEN) syndrome inherited in an autosomal-dominant manner and characterized by cardiac and noncardiac myxomatous tumors, multiple endocrine tumors, and distinctive pigmented lesions of the skin.^17^ According to the National Organization for Rare Diseases (NORD), approximately 600 affected individuals have been reported since the disorder was first described in the medical literature in 1985.^18^ Inactivating mutations in the Protein Kinase cAMP-Dependent Type I Regulatory Subunit Alpha (PRKAR1A) protein are found in approximately 70% of CNC cases and are closely associated with other MEN syndromes (17-19).^19,20^ Thus, we sought to identify subgraph patterns between existing drugs and Carney Complex that required genetic associations with either PRKAR1A or other genes in the pathway.

Exploration of the DD4 subgraph (see Figure 2B) involving CNC and PRKAR1A was not possible, as there are no on-market drugs with activity against PRKAR1A that could be found in either the public domain or internally to AbbVie. Thus, we expanded to other genes that share a pathway with PRKAR1A as defined by the Consensus Path Database (CPDB) in order to construct potential DD5 subgraphs (see Figure 2C). With this expansion, a directed DD5 subgraph pattern for drug repurposing for CNC was identified, as illustrated in Figure 3.

**Figure 2.**
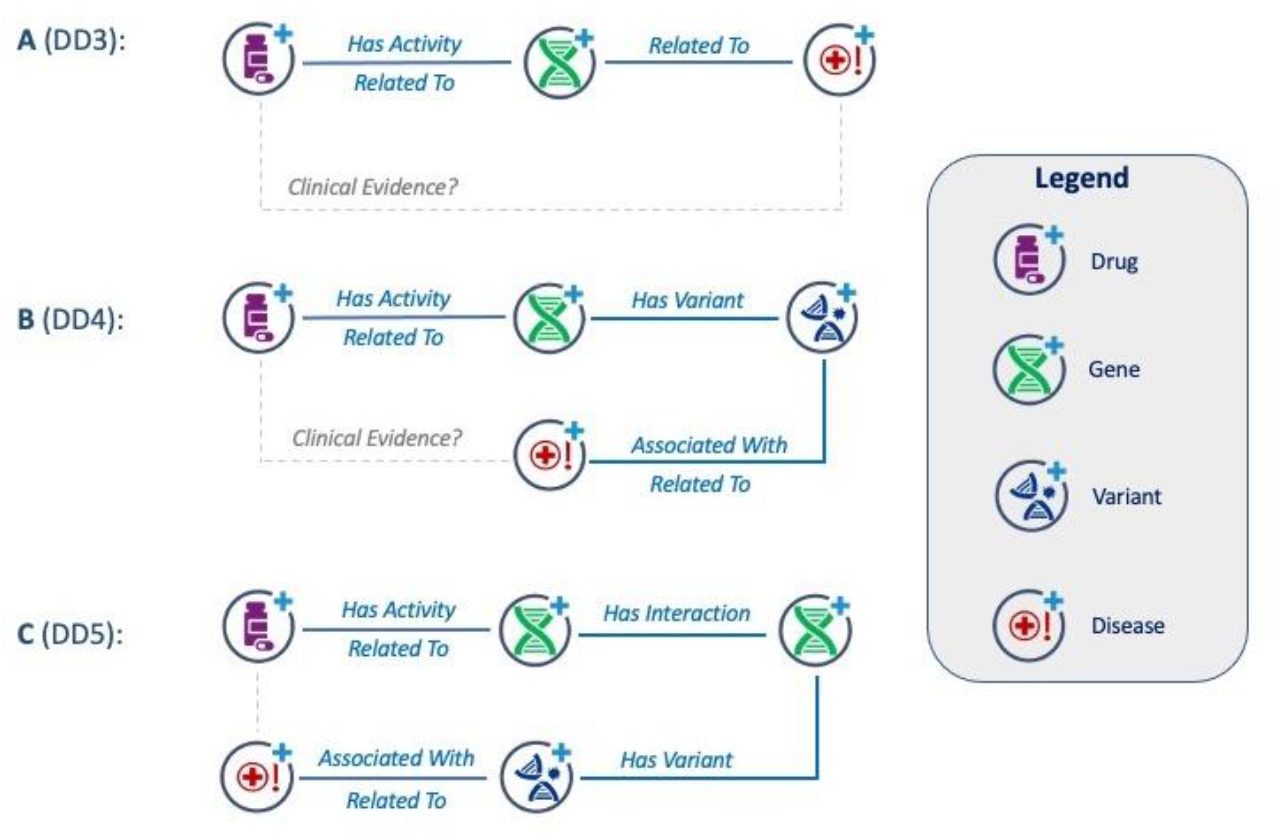
Drug-Disease subgraphs of A) length 3 (DD3), B) length 4 (DD4), and length 5 (DD5) as described in the text. Subgraphs were validated using clinical evidence to support a therapeutically relevant Drug-Disease association as shown in Table 1.

**Figure 3.**
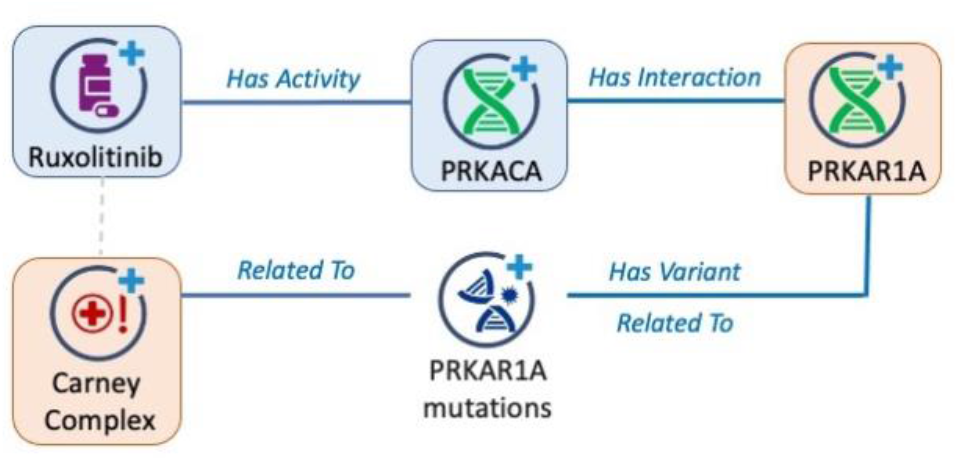
Directed DD5 subgraph for identification of drug repurposing opportunities for CNC. The subgraph was required to contain both PRKAR1A and CNC, as highlighted in orange. Ruxilitinib, which exhibits activity against PRKACA, was identified as a potential option.

As shown in Table 2, there are 27 genes that share a pathway with PRKAR1A at the highest confidence level (CPDB confidence score > 0.99). Of these genes, only GSK2B, MTOR, PRKACA, PRKACB, and RPS17 have on-market drugs with at least moderate (IC_50_ < 1 μM) reported activity. It is important to note that for many of these drugs, the reported activity against the gene is not the primary or designed mechanism of action.

**Table 2.** Genes that share a pathway with PRKAR1A, with availability of on-market drugs with at least moderate (sub-μM) potency.

| Gene | On-Market Drug? ($IC_{50} < 1 \mu M$) |
| --- | --- |
| GSK3B | Yes |
| MTOR | Yes |
| PRKACA | Yes |
| PRKACB | Yes |
| RPS17 | Yes |
| AGO2 | None found* |
| AGO3 | None found |
| AKAP1 | None found |
| AKAP4 | None found |
| BCL2L11 | None found |
| BTRC | None found |
| CCNE1 | None found |
| CEP250 | None found |
| CUL3 | None found |
| CUL7 | None found |
| GRB2 | None found* |
| MEN1 | None found* |
| MYO7A | None found |
| PPP1R12A | None found |
| PRKACG | None found |
| PRKAR1B | None found |
| PRKAR2B | None found |
| PYCARD | None found |
| SMAD2 | None found |
| SOX2 | None found |
| TAB1 | None found |
| YWHAZ | None found |

\*Only weak ( $IC_{50} > 1 \mu M$ ) inhibitors could be found against these genes

Review of the literature evidence in the subgraph analysis revealed PRKACA to be of particular interest in connection to PRKAR1A. PRKAR1A is a regulatory protein, and the inactivating mutations in PRKAR1A disrupt the interaction with PRKACA and lead to increased catalytic activity of the PRKACA oncogene.^19^ Over 125 pathogenic PRKAR1A mutations have been identified to date in CNC cases, caused by single base pair substitutions, small insertions/deletions (<15bp), combined rearrangements, and large deletions. Functionally, these mutations result in increased PKA-dependent cAMP signaling.^21^ Mutations in PRKACA that disrupt binding to the PRKAR1A regulatory domain also increase-PKA-dependent cAMP signaling and are found in patients with Cushing Syndrome and other adrenocortical disorders. Both of these pieces of genetic evidence support the potential of PRKACA inhibitors in the treatment of CNC and other forms of hypercortisolism, and in fact the use of small molecule PRKACA inhibitors in this disease setting was postulated as far back as 2014.^22^ In 2015, Berthon et al. reported that inactivation of PRKAR1A leads to constitutive activation of PRKACA and adrenocortical, suggesting that inactivation of PRKACA may be a means to treat CNC.^23^

There are no on-market drugs have been developed against PRKACA as the primary mechanism of action. However, the subgraph analysis captured the reported off-target activity of Ruxolitinib (Incyte, trade name Jakafi) against PRKACA as reported by the HMS LINCS Project^24^ and Eberl et al.^25^ It is important to note that target activity alone is insufficient to quality a drug for repurposing, and many other considerations must be evaluated to understand therapeutic potential, including drug exposure and pharmacokinetic data, efficacy in current indications, human safety profiles, and more. A key consideration is whether drug exposures achieved at approved doses are sufficiently high to achieve a therapeutic effect relative to PRKACA activity, especially since this off-target activity is likely significantly reduced relative to the reported mechanism of action for the drug.

We illustrate some of these considerations for Ruxolitinib, a JAK1/2 inhibitor approved for the treatment of myelofibrosis. According to the cellular activity shown in Table 3, Ruxolitinib is approximately 10- to 100-fold less potent against PRKACA than JAK1/2 (the primary mechanism of action for the approved indications).^26^ At the approved dose of 20 mg, the observed C_max_ level for Ruxolitinib in healthy volunteers is between 650 and 1160 nM.^25^ This results in a C_max_/IC_50_ ratio of ~6 for PRKACA, suggesting that at least some level of PRKACA inhibition may be achieved with Ruxolitinib at currently approved oral doses. However, this ratio is substantially lower than that achieved for JAK1 and Jak2 (C_max_/IC_50_ ratio > 300 in both cases, see Table 4). While this is lower target coverage of PRKACA relative to JAK1/2, it is not known what level (or duration) of PRKACA inhibition may be required to restore normal PKA-dependent cAMP signaling. It is also not known what role JAK1/2 inhibition may play in MEN disorders. The fact that Ruxolitinib is approved in an oncology setting makes it an intriguing opportunity for further clinical and non-clinical exploration in Carney Complex and other MEN disorders.

**Table 3.** Potency and estimated exposures of Ruxolitinib, against Jak1/2 and PRKACA

| Target | Ruxolitinib |  |
| --- | --- | --- |
| | Cellular Potency (nM)<sup>a</sup> | Human $C_{max}/IC_{50}$<sup>b</sup> |
| JAK1 | 1 | 1000 |
| JAK2 | 3 | 333 |
| PRKACA | 160 | 6.3 |

<sup>a</sup>Taken from Eberl et al.<sup>25</sup>
<sup>b</sup>Based on Human $C_{max}$ of 1000 nM reported for recommended 20 mg/day dose of Ruxolitinib from Shi et al.<sup>26</sup>

## Conclusion

In this paper, we have demonstrated the potential behind utilizing a 911M edge knowledge graph built from publicly available data sources for drug repurposing. Of particular note is the utilization of NLP to mine unstructured biomedical text to enrich the knowledge graph with connections and evidence that cannot be derived strictly from structured sources. We leverage subgraph analysis against the knowledge graph to calculate all possible genetically supported Drug-Gene-Disease paths of length 4 (DD4) and length 5 (DD5, expanded by pathway membership). Consistent with other studies, these genetically supported subgraphs show significant enrichment of clinically validated drug-indication pairs, supporting their use in disease expansion and drug repurposing initiatives.

We have also illustrated the application of such subgraphs in the case of drug repurposing for CNC, a rare genetic disease for which no current therapies exist. While no drugs have been designed against PRKACA, chemical proteomic studies revealed moderate drug activity against (see Tables 3 and 4), and this information is captured and exposed as connections in the graph. Further exploration of the human pharmacokinetic profile of Ruxolitinib (developed as a JAK1/2 inhibitor) indicates that at least some level of PRKACA inhibition is achieved at currently approved doses. As a criticism, we have no evidence that Ruxolitinib will work for CNC. However, an on-market drug with a reasonable safety profile and with a label for cancer (myelofibrosis in the case of Ruxolitinib) could be considered for patients with CNC. While we recognize this is a critical decision between the prescribing physician and patient, it is also important to note that many rare diseases will not be large enough to conduct a clinical trial. We hope our strategy here in utilizing subgraph analysis derived from compiled knowledge graphs could be leveraged by the Pharmaceutical Industry in a pre-competitive forum in partnership with payers, providers, and patients. Though not directly utilized in this analysis, we believe that additional use of our causal expression curation to enrich the knowledge graph with cause-effect relationships can be used to expand this subgraph analysis strategy to a range of applications beyond drug repurposing. We encourage developing a series of pilots to evaluate this opportunity.

## Supporting information

Drug-Indication Pairs Used for Subgraph Analysis

## References

1. Wouters, O. J., McKee, M. & Luyten, J. Estimated Research and Development Investment Needed to Bring a New Medicine to Market, 2009-2018. Jama 323, 844–853 (2020).

2. Shahreza, M. L., Ghadiri, N., Mousavi, S. R., Varshosaz, J. & Green, J. R. A review of network-based approaches to drug repositioning. Brief Bioinform 19, 878–892 (2017).

3. Wong, C. H., Siah, K. W. & Lo, A. W. Estimation of clinical trial success rates and related parameters. Biostatistics 20, 273–286 (2019).

4. Naylor, S. & JM, S. Therapeutic Drug Repurposing, Repositioning and Rescue Part I: Overview - Drug Discovery World (DDW). https://www.ddw-online.com/therapeutic-drug-repurposing-repositioning-and-rescue-part-i-overview-1463-201412/ (2014).

5. Cha, Y. et al. Drug repurposing from the perspective of pharmaceutical companies. Brit J Pharmacol 175, 168–180 (2018).

6. Wu, Z., Wang, Y. & Chen, L. Network-based drug repositioning. Mol Biosyst 9, 1268–1281 (2013).

7. Cheng, F. et al. Network-based approach to prediction and population-based validation of in silico drug repurposing. Nat Commun 9, 2691 (2018).

8. Gysi, D. M. et al. Network medicine framework for identifying drug-repurposing opportunities for COVID-19. Proc National Acad Sci 118, e2025581118 (2021).

9. Mullen, J., Cockell, S. J., Tipney, H., Woollard, P. M. & Wipat, A. Mining integrated semantic networks for drug repositioning opportunities. Peerj 4, e1558 (2016).

10. Schultz, B. et al. A method for the rational selection of drug repurposing candidates from multimodal knowledge harmonization. Sci Rep-uk 11, 11049 (2021).

11. Subramanian S., Baldini I., Ravichandran S., Katz-Rogozhnikov D.A., Natesan Ramamurthy K., Sattigeri P., Varshney K.R., Wang A., Mangalath P., & Kleiman L.B. A natural language processing system for extracting evidence of drug repurposing from scientific publications. Proceedings of the AAAI Conference on Artificial Intelligence 34, 13369–13381 (2020).

12. Öztürk H., Özgür A., Schwaller P., Laino T., & Ozkirimli E. Exploring chemical space using natural language processing methodologies for drug discovery. Drug Discovery Today 25, 689–705 (2020).

13. Kong H.J. Managing Unstructured Big Data in Healthcare System. Healthcare Informatics Research 25, 1–2 (2019).

14. King, E. A., Davis, J. W. & Degner, J. F. Are drug targets with genetic support twice as likely to be approved? Revised estimates of the impact of genetic support for drug mechanisms on the probability of drug approval. Plos Genet 15, e1008489 (2019).

15. Nelson, M. R. et al. The support of human genetic evidence for approved drug indications. Nat Genet 47, 856–860 (2015).

16. Roessler, H. I., Knoers, N. V. A. M., Haelst, M. M. van & Haaften, G. van. Drug Repurposing for Rare Diseases. Trends Pharmacol Sci 42, 255–267 (2021).

17. Courcoutsakis N.A., Tatsi C., Patronas N.J., Lee C.C.R., Prassopoulos P.K., & Stratakis C.A. The complex of myxomas, spotty skin pigmentation and endocrine overactivity (Carney complex): imaging findings with clinical and pathological correlation. Insights Into Imaging 4, 119–33 (2013).

18. Carney Complex - NORD (National Organization for Rare Disorders). https://rarediseases.org/rare-diseases/carney-complex/#:~:text=Carney%20complex%20is%20a%20rare,the%20skin%20of%20affected%20areas.

19. Bossis, I. & Stratakis, C. A. Minireview: PRKAR1A: Normal and Abnormal Functions. Endocrinology 145, 5452–5458 (2004).

20. Papanastasiou, L. et al. Identification of a novel mutation of the PRKAR1A gene in a patient with Carney complex with significant osteoporosis and recurrent fractures. Hormones 15, 129–35 (2015).

21. Liu, Q. et al. Carney complex with PRKAR1A gene mutation: A case report and literature review. Medicine 6, e8999 (2017).

22. Stratakis, C. A. E Pluribus Unum? The Main Protein Kinase A Catalytic Subunit (PRKACA), a Likely Oncogene, and Cortisol-Producing Tumors. J Clin Endocrinol Metabolism 99, 3629–3633 (2014).

23. Berthon, A. S., Szarek, E. & Stratakis, C. A. PRKACA: the catalytic subunit of protein kinase A and adrenocortical tumors. Frontiers Cell Dev Biology 3, 26 (2015).

24. Gaulton, A. et al. ChEMBL: a large-scale bioactivity database for drug discovery. Nucleic Acids Res 40, D1100–D1107 (2012).

25. Eberl, H. C. et al. Chemical proteomics reveals target selectivity of clinical Jak inhibitors in human primary cells. Sci Rep-uk 9, 14159 (2019).

26. Shi, J. G. et al. The Pharmacokinetics, Pharmacodynamics, and Safety of Orally Dosed INCB018424 Phosphate in Healthy Volunteers. J Clin Pharmacol 51, 1644–1654 (2011).

