## Supplemental Tables for "Leveraging a Billion-Edge Knowledge Graph for Drug Re-purposing and Target Prioritization using Genomically-Informed Subgraphs"

**Appendix A.** Knowledge graph data sources and relationship types

| **Source(s)** | **Entity 1** | **Relationship** | **Entity 2** |
|---|---|---|---|
| **Knowledge from Structured Sources** | | | |
| HMS LINCS | Drug | *has_activity* | Gene |
| dbSNP | Gene | *has_variant* | Variant |
| | Variant | *associated_with* | Disease |
| HGNC | Gene | *is_member_of* | Gene (Family) |
| CPDB | Gene | *has_interaction* | Gene |
| GWAS Catalog | Gene | *related_to* | Phenotype |
| | Gene | | Disease |
| | Variant | | Phenotype |
| | Disease | | Variant |
| **Knowledge from Unstructured Sources** | | | |
| PubMed<br>PMC<br>bioRxiv<br>medRxiv<br>arXiv<br>Google Patents<br>ClinicalTrials.gov | Gene | *related_to* | Gene |
| | Gene | | Disease |
| | Gene | | Phenotype |
| | Gene | | Variant |
| | Gene | | Cell Type |
| | Gene | | Drug |
| | Disease | | Disease |
| | Disease | | Phenotype |
| | Disease | | Variant |
| | Disease | | Cell Type |
| | Disease | | Drug |
| | Drug | | Variant |
| | Drug | | Cell Type |
| | Drug | | Phenotype |
| | Phenotype | | Cell Type |
| | Phenotype | | Variant |
| | Variant | | Cell Type |

**Table A1.** To construct the knowledge graph, structured and unstructured data sources used for relationship extraction. Entity pair relationship types derived from each source is listed above.

©2022 TELLIC LLC

**Appendix B**. Increasing Clinical Support to Phase 2+ vs Phase 1+

| Subgraph Pattern | Total Subgraph Counts | Drug-Indication Subgraph Counts[a] | Documented Phase 2+ | Clinically Supported | Enrichment |
|---|---|---|---|---|---|
| All Pairs | 15,773,967 | 15,773,967 | 82,076 | 0.52% | |
| DD3 | 36,101,662 | 4,429,489 | 62,514 | 1.41% | 2.71 |
| DD4-1 (≥1 variant) | 505,087[b] | 212,219 | 16,658 | 7.85% | 15.09 |
| DD4-2 (≥2 variants) | 177,335[b] | 90,246 | 10,367 | 11.49% | 22.08 |
| DD4-3 (≥3 variants) | 78,252[b] | 47,464 | 6,836 | 14.40% | 27.68 |
| DD4-4 (≥4 variants) | 47,300[b] | 30,963 | 4,844 | 15.64% | 30.07 |
| DD4-5 (≥5 variants) | 30,570[b] | 20,618 | 3,628 | 17.60% | 33.82 |
| DD5-1 (≥1 variant) | 4,277,013[b] | 732,749 | 26,629 | 3.63 | 6.98 |
| DD5-2 (≥2 variants) | 1,654,709[b] | 358,475 | 18,206 | 5.08 | 9.76 |
| DD5-3 (≥3 variants) | 906,071[b] | 228,383 | 13,758 | 6.02 | 11.58 |
| DD5-4 (≥4 variants) | 638,230[b] | 170,503 | 10,883 | 6.38 | 12.27 |
| DD5-5 (≥5 variants) | 499,600[b] | 135,651 | 8,961 | 6.61 | 12.70 |

[a]Subgraph limited to the 1749 Diseases treated (in Phase 2+) by the set of 5767 drugs used in the analysis
[b]Unique Drug-Gene-Disease paths regardless of number of intervening variants

**Table B1.** For each subgraph type, the proportion of subgraph counts with clinical support and subsequent enrichment compared to “All Pairs” were recalculated based on documented phase 2 (or later) clinical trials.

**Appendix C**. Separating Oncology and Non-Oncology Indications

| Subgraph Pattern | Total Subgraph Counts | Drug-Indication Subgraph Counts[a] | Documented Phase 1+ | Clinically Supported | Enrichment |
|---|---|---|---|---|---|
| All Pairs | 2,994,706 | 2,994,706 | 34,601 | 1.16% | |
| DD3 | 8,783,084 | 878,463 | 27,214 | 3.10% | 2.68 |
| DD4-1 (≥1 variant) | 160,184[b] | 57,574 | 7,863 | 13.66% | 11.82 |
| DD4-2 (≥2 variants) | 71,496[b] | 32,400 | 5,531 | 17.07% | 14.77 |
| DD4-3 (≥3 variants) | 39,643[b] | 22,302 | 4,190 | 18.79% | 16.26 |
| DD4-4 (≥4 variants) | 26,656[b] | 17,276 | 3,400 | 19.68% | 17.03 |
| DD4-5 (≥5 variants) | 19,317[b] | 13,452 | 2,902 | 21.57% | 18.67 |
| DD5-1 (≥1 variant) | 1,625,749[b] | 207,148 | 13,895 | 6.71 | 5.81 |
| DD5-2 (≥2 variants) | 674,699[b] | 132,061 | 10,604 | 8.03 | 6.95 |
| DD5-3 (≥3 variants) | 380,429[b] | 105,296 | 9,009 | 8.56 | 7.41 |
| DD5-4 (≥4 variants) | 270,726[b] | 87,251 | 7,638 | 8.75 | 7.58 |
| DD5-5 (≥5 variants) | 221,708[b] | 78,141 | 7,047 | 9.02 | 7.81 |

[a]Subgraph limited to the 481 oncology indications treated by the set of 6226 drugs used in the analysis
[b]Unique Drug-Gene-Disease paths regardless of number of intervening variants

**Table C1.** Subgraph pattern counts were calculated for oncology indications only, along with the subset of patterns supported by Phase 1+ clinical evidence for DD3 through DD5 patterns.

| Subgraph Pattern | Total Subgraph Counts | Drug-Indication Subgraph Counts[a] | Documented Phase 1+ | Clinically Supported | Enrichment |
|---|---|---|---|---|---|
| All Pairs | 14,687,134 | 14,687,134 | 62,106 | 0.42% | |
| DD3 | 28,521,851 | 3,830,000 | 46,211 | 1.21% | 2.85 |
| DD4-1 (≥1 variant) | 356,860[b] | 161,426 | 11,322 | 7.01% | 16.59 |
| DD4-2 (≥2 variants) | 110,420[b] | 60,812 | 6,362 | 10.46% | 24.74 |
| DD4-3 (≥3 variants) | 40,837[b] | 26,793 | 3,662 | 13.67% | 32.32 |
| DD4-4 (≥4 variants) | 22,140[b] | 14,824 | 2,163 | 14.59% | 34.51 |
| DD4-5 (≥5 variants) | 12,384[b] | 8,035 | 1,293 | 16.09% | 38.06 |
| DD5-1 (≥1 variant) | 2,841,758[b] | 563,371 | 17,407 | 3.09 | 7.31 |
| DD5-2 (≥2 variants) | 1,055,388[b] | 244,522 | 10,844 | 4.43 | 10.49 |
| DD5-3 (≥3 variants) | 567,769[b] | 134,738 | 7,266 | 5.39 | 12.75 |
| DD5-4 (≥4 variants) | 397,552[b] | 92,065 | 5,255 | 5.71 | 13.50 |
| DD5-5 (≥5 variants) | 301,768[b] | 64,701 | 3,665 | 5.66 | 13.40 |

[a]Subgraph limited to the 2359 non-oncology indications treated by the set of 6226 drugs used in the analysis
[b]Unique Drug-Gene-Disease paths regardless of number of intervening variants

**Table C2.** Subgraph pattern counts were calculated for non-oncology indications only, along with the subset of patterns supported by Phase 1+ clinical evidence for DD3 through DD5 patterns.

©2022 TELLIC LLC
